# Integrative Statistical Inferences for Drug Sensitivity Biomarkers in Cancer

**DOI:** 10.1101/194670

**Authors:** Ehsan Ullah, Saila Shama, Noora Al Muftah, Ian Richard Thompson, Reda Rawi, Raghvendra Mall, Halima Bensmail

**Affiliations:** Qatar Computing Research Institute, Hamad Bin Khalifa University, Doha, Qatar; Faculty of Applied Science and Engineering, Engineering Science, University of Toronto, 40 Saint George Street, Toronto, Canada M4C 3C2; Department of Biostatistics, Harvard School of Public Health, Boston, MA 02115, USA; Qatar Biomedical Research Institute, Hamad Bin Khalifa University, Doha, Qatar; Vaccine Research Center, National Institute of Allergy and Infectious Diseases, National Institutes of Health,Bethesda, MD 20814, USA

## Abstract

Personal medicine has been associated with different patient responses to different anti-cancer therapies. Recently, scientists are looking not only for new biomarkers associated with a disease such as cancer but also identifying biomarkers that predict patients who are most likely to respond to a particular cancer treatment. Orderly endeavors to relate cancer mutational information with biological conditions may encourage the interpretation of somatic mutation indexes into significant biomarkers for patient stratification.

We have screened and incorporated a board of cancer cell lines from Genomics of Drug Sensitivity in Cancer (GDSC) database to recognize genomic highlights related with drug sensitivity. We used mutation, DNA copy number variation, and gene expression information from Catalogue of Somatic Mutations in Cancer (COSMIC) and The Cancer Genome ATLAS (TCGA) for cell lines with their reactions to associate focused and cytotoxic treatments with approved drugs and drugs under clinical and preclinical examination.

We discovered mutated cancer genes were related with cell reaction to, mostly accessible, cancer medications and some mutated genes were related with sensitivity to an expansive scope of therapeutic agents. By connecting drug activity to the useful many-sided quality of cancer genomes, efficient pharmacogenomic profiling in tumor cell lines gives an intense biomarker revelation stage to guide balanced malignancy remedial systems.

Our study highlights that gene ANK2 amplification, and gene CELSER1 amplification and deletion are highly associated with anti-leukemic drug candidate LFM-A13. It also highlights that gene NUP214 and ROS1 copy number and gene NSD1 amplification are as a group highly associated with the parkinson drug Nilotinib. Finally, our study confirms that gene BRAF mutation is interacting with the BRAF-selective inhibitors drugs PLX4720 and SB590885. On the other hand, our study provides two open source analysis packages: bastah for the multitask-association analysis, and UNGeneAnno for automatic annotation of the variants.

## 1 Background

Clinical reactions to anti-cancer treatments are regularly confined to a subset of patients. Now and again, mutated cancer genes are strong biomarkers of reaction to targeted agents. There is an earnest need to recognize biomarkers that can anticipate patients ability to react to a treatment [32]. Moreover, the success of precision medicine is based on our ability to effectively translate genomic data into effective, customized prognosis and treatment strategies for individual patients. This requires identifying a genomic disease signature from a patient, then associating it with the most effective therapeutic intervention [6]. Recently, researchers have utilized cell lines to identify molecular consequences associated with various exposures, such as drugs, small molecules, or pathogens to move towards this goal [13], but many challenges still remain including what kind of data is needed to develop these genomic signatures and what kind of methods are needed to extract the appropriate information from high-throughput genomic data sets. Such analysis requires the use of genomic features such as gene expression, copy number variation, deletion, mutation, and sequence data, in the predictive modeling of anti-cancer drug response and holds the potential to speed up the emergence of personalized cancer therapies.

The first venture to address these challenges is to create far reaching drug affectability profiling estimations crosswise over different drugs, malady sub-sorts, and genomic profiling advances. Several of these data sets have been generated with a focus on cancer biology, in particular Glioma and Breast cancer. The bottleneck then becomes robust computational approaches that connect genomic profiles with drug and disease response in these datasets. For example, we view each tumor’s Copy Number Aberration (CNA) and mutation profiles as a system perturbation that simultaneously affect multiple genes. CNAs and mutations tend to appear in a patient-specific and multifactorial manner that is ideal for targeted medicine and targeted drug sensitivity [12].

In this paper, we present SLICED (**S**tatistica**L I**nferences for **CE**ll lines screening in **D**rug discovery), a computational method that links genomic features, drugs and the associated half maximal inhibitory concentration (IC50) i.e., concentration at which half the target cells are killed. SLICED uses a newly developed two-layer algorithm which performs a multiplex association analysis on genomic features drugs cell lines with a focus on lasso (least absolute shrinkage and selection operator) [25].

We identified clinical trial datasets that had assessed tumor gene expression before drug treatment (using expression microarrays, copy numbers and mutations) and had subsequently measured a clear drug response phenotype using The Cancer Genome Atlas (TCGA), Catalogue of Somatic Mutations in Cancer (COSMIC) and Genomics of Drug Sensitivity in Cancer (GDSC) consortium to which we apply SLICED to predict the association of drugs-genomic features.

## 2 Results

Our goal was to use baseline gene expression and in vitro drug sensitivity derived from cell lines, coupled with in vivo baseline tumor gene expression, mutation and copy number to predict patients response to anti-cancer drugs. An overview of our approach is shown in Figure 1 (complete details are given in Materials and Methods section).

**Figure 1:**
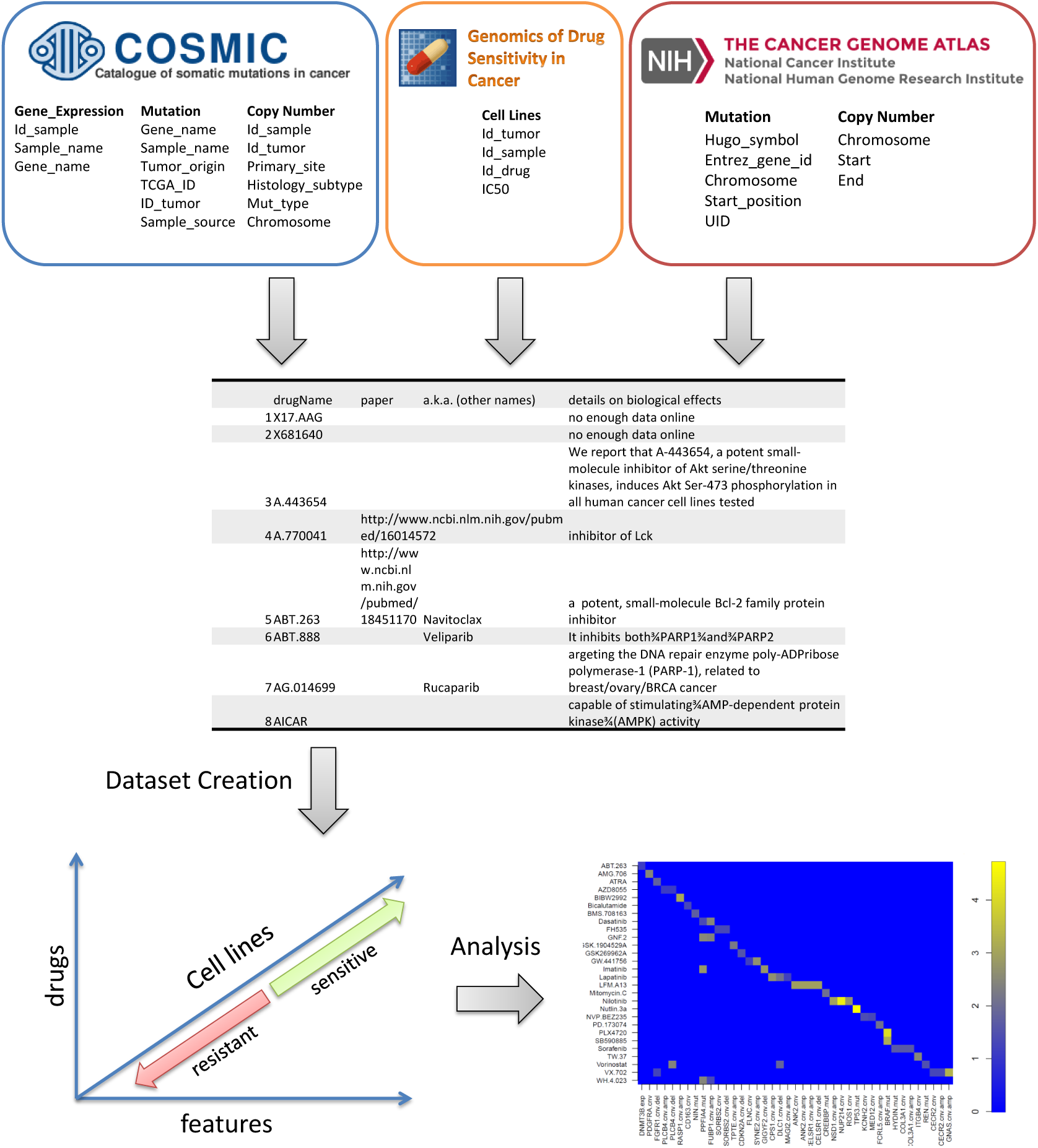
Flowchart of SLICED.

To capture the high degree of genomic diversity in cancer and to identify rare mutant subsets with altered drug sensitivity, we assembled 639 human tumour cell lines, representing the spectrum of common and rare types of adult and childhood cancers of epithelial, mesenchymal and haematopoietic origin. Cell lines were subjected to sequencing of the full coding exons of 64 commonly mutated cancer genes, genome-wide analysis of copy number gain and loss using Affymetrix SNP6.0 microarrays, and expression profiling using Affymetrix HT-U133A microarrays.

The available drugs covered a wide range of targets and processes implicated in cancer biology. They encompassed both targeted agents and cytotoxic chemotherapeutics, including approved drugs used in clinical practice, drugs in development undergoing studies in clinical trials, and experimental tool compounds (list of drugs is available as supplementary material S1 and annotations as S2). To dig into drug-to-drug variation, we included multiple drugs (related to the cancer) designed against well-credentialed targets. The effect of 72 hours of drug treatment on cell viability was used to derive a multi-parameter description of drug sensitivity (IC50), and the slope of the dose response curve. In preliminarily tests, we assessed several of the plethora of available machine learning algorithms, including principal component regression [16], elastic net regression [36], sslasso [14] and lasso.proj [27]. Among them, elastic net, sslasso and lasso.proj were consistently the best performers, with the added advantage of having an elasticNet being highly computationally efficient which is crucial for cross-validation analysis, but does not produce the significance of the effect of the association (p-value). Sslasso and lasso.proj which provide significance of the effect (p-value) are unfortunately computationally inefficient.

In the combined population, 1025 gene expressions, 1091 mutation and 476 CNV were evaluated. Using 

~~~
bastah
~~~

, we find 41 genomic features whose expression significantly correlated to IC50 and 28 drugs (see Table 1). We identified mutations that were significantly associated with IC50 through regulation, gene expressions and deletion. In the context of compound sensitivity prediction, previous studies have demonstrated that such feature selection may provide the basis for identifying functional biomarkers underlying compound sensitivity or resistance [21, 9, 3, 11].

**Table 1:** Final list of significant association between drugs and genomics features using bastah

| Drug | Approved | Feature | p-val | $\hat{\beta}$ |
| --- | --- | --- | --- | --- |
| Nutlin-3a | | TP53.mut | $3.63 \times 10^{-49}$ | 2.27 |
| Nilotinib (Tasigna) | yes | NUP214.cnv | $1.31 \times 10^{-36}$ | -1.15 |
| PLX-4720 | | BRAF.mut | $1.63 \times 10^{-25}$ | -1.09 |
| VX-702 | | GNAS.cnv.amp | $2.93 \times 10^{-12}$ | 0.44 |
| SB590885 | | BRAF.mut | $3.88 \times 10^{-11}$ | -0.72 |
| BIBW2992 (Afatinib) | yes | GPRASP1.cnv.amp | $2.53 \times 10^{-10}$ | -0.63 |
| Nilotinib (Tasigna) | yes | NSD1.cnv.amp | $2.11 \times 10^{-09}$ | -0.62 |
| LFM-A13 | | CELSR1.cnv.del | $5.64 \times 10^{-09}$ | 0.36 |
| LFM-A13 | | ANK2.cnv.amp | $6.19 \times 10^{-09}$ | 0.36 |
| LFM-A13 | | CELSR1.cnv.amp | $6.80 \times 10^{-09}$ | 0.36 |
| Lapatinib | yes | MAGI2.cnv.amp | $7.63 \times 10^{-09}$ | -0.61 |
| Imatinib (STI-571) | yes | PPFIA4.mut | $6.35 \times 10^{-08}$ | -0.94 |
| Imatinib (STI-571) | yes | GIGYF2.cnv.del | $6.66 \times 10^{-08}$ | -0.92 |
| Nilotinib (Tasigna) | yes | ROS1.cnv | $2.35 \times 10^{-07}$ | -0.57 |
| GW-441756 | | SYNE2.cnv.amp | $3.90 \times 10^{-07}$ | -0.42 |
| Lapatinib | yes | CPS1.cnv.amp | $5.05 \times 10^{-07}$ | -0.90 |
| GNF-2 | | PPFIA4.mut | $4.10 \times 10^{-06}$ | -0.74 |
| Dasatinib | yes | FUBP1.cnv.amp | $5.03 \times 10^{-06}$ | -1.96 |
| TW-37 | | ITGB4.cnv | $5.62 \times 10^{-06}$ | 0.41 |
| WH.4.023 | | PPFIA4.mut | $8.00 \times 10^{-06}$ | -2.11 |
| AMG-706 (Motesanib) | | PDGFRA.cnv | $8.32 \times 10^{-06}$ | -0.38 |
| GNF-2 | | FUBP1.cnv.amp | $1.28 \times 10^{-05}$ | -0.69 |
| Vorinostat | yes | PLCB4.cnv.del | $1.28 \times 10^{-05}$ | 0.35 |
| GSK-1904529A | | TPTE.cnv.amp | $7.13 \times 10^{-05}$ | -0.31 |
| Lapatinib | yes | DLC1.cnv.del | $2.49 \times 10^{-04}$ | -0.75 |
| PD173074 | | FCRL5.cnv.amp | $3.44 \times 10^{-04}$ | -0.41 |
| Mitomycin-C | yes | CREBBP.mut | $3.68 \times 10^{-04}$ | -0.66 |
| Vorinostat | yes | DLC1.cnv.del | $1.88 \times 10^{-03}$ | 0.30 |
| ATRA (Tretinoin) | | FGFR1.cnv.del | $2.46 \times 10^{-03}$ | -0.41 |
| BMS-708163 (Avagacestat) | | NIN.mut | $3.29 \times 10^{-03}$ | -0.31 |
| Sorafenib | yes | HYDIN.mut | $5.64 \times 10^{-03}$ | -0.73 |
| Sorafenib | yes | COL3A1.cnv.amp | $5.90 \times 10^{-03}$ | -0.73 |
| Sorafenib | yes | COL3A1.cnv | $6.07 \times 10^{-03}$ | -0.73 |
| GSK269962A | | CDKN2A.cnv.del | $8.91 \times 10^{-03}$ | -0.84 |
| Vorinostat | yes | REN.mut | $1.24 \times 10^{-02}$ | 0.28 |
| NVP-BEZ235 | | KCNH2.cnv | $1.51 \times 10^{-02}$ | -0.36 |
| Dasatinib | yes | PPFIA4.mut | $1.91 \times 10^{-02}$ | -1.50 |
| WH.4.023 | | FUBP1.cnv.amp | $2.18 \times 10^{-02}$ | -1.54 |
| FH535 | | SORBS2.cnv.del | $2.21 \times 10^{-02}$ | -0.38 |
| FH535 | | SORBS2.cnv | $2.28 \times 10^{-02}$ | -0.38 |
| Bicalutamide | yes | CD163.cnv | $2.32 \times 10^{-02}$ | -0.14 |
| NVP-BEZ235 | | MED12.cnv | $2.43 \times 10^{-02}$ | 0.35 |
| VX-702 | | CECR2.cnv.amp | $2.54 \times 10^{-02}$ | -0.24 |
| VX-702 | | CECR2.cnv | $2.60 \times 10^{-02}$ | -0.24 |
| VX-702 | | FGFR1.cnv.del | $2.85 \times 10^{-02}$ | -0.24 |
| LFM-A13 | | ANK2.cnv | $3.67 \times 10^{-02}$ | 0.36 |
| GW-441756 | | FLNC.cnv | $3.95 \times 10^{-02}$ | -0.29 |
| ABT-263 (Navitoclax) | | DNMT3B.exp | $4.63 \times 10^{-02}$ | -1.00 |
| AZD8055 | | PLCB4.cnv.del | $4.70 \times 10^{-02}$ | 0.28 |
| AZD8055 | | PLCB4.cnv.amp | $4.75 \times 10^{-02}$ | 0.28 |

We compiled a list of known biomarkers of sensitivity (gold standards) for compounds represented in both the Sanger and CCLE panels, cell lines were categorized into 13 tissues, and evaluated the rank of each biomarker (based on the absolute value of the regression coefficients) in the model generated by bastah for the corresponding compound. In the following, we summarize the most important significant associations identified by our algorithm with a p-value threshold of 10^-06^. In particular we focus on two categories of drugs: approved and under clinical trials.

### 2.1 Association of biomarker sensitivity with approved drugs

- Nilotinib (associated with NSD1 amplification, and *NUP214* and *ROS1* copy number change): Nilotinib is a drug used for chronic myelogenous leukaemia (CML) [30].
- Afatinib (associated with GPRASP1 amplification): Afatinib is a tyrosine kinase inhibitor of human EGF receptor (EGFR) and human epidermal growth factor receptor-2 (HER-2)/neu. EGFR and HER-2/neu used for treatment of solid tumors [18].
- Lapatinib (associated with CPS1 and MAGI2 amplification): Lapatinib is a HER2-positive breast cancer medicine. It is a dual tyrosine kinase inhibitor that interrupts the epidermal growth factor receptor (EGFR) and HER2/neu pathways [35].
- Imatinib (associated with GIGYF2 deletion and PPFIA4 mutation): Imatinib is a tyrosine-kinase inhibitor. It is used in the treatment of multiple cancers, most notably Philadelphia chromosome-positive (Ph+) chronic myelogenous leukemia (CML) [4].
- Dasatinib (associated with PPFIA4 mutation): Dasatinib is an inhibitor of BCR-ABL tyrosine kinase and SRC family tyrosine kinase used to treat patients with chronic myelogenous leukemia (CML) [20].

### 2.2 Association of biomarker sensitivity with under clinical trial drugs

- Nutlin-3 (associated with *TP53* mutation): Nutlin-3 is a potent inhibitor of MDM2 (mouse double minute 2) binding to p53 that induces the expression of p53-regulated genes [5].
- PLX-4720 (associated with *BRAF* mutation): PLX-4720 significantly inhibits the ERK phosphorylation in cell lines bearing B-Raf^V^600^E^, but not the cells with wild-type B-Raf. PLX-4720 significantly inhibits the growth of tumor cell lines bearing the B-Raf^V^600^E^ oncogene [22].
- VX-702 (associated with *GNAS* amplification): VX-702 is an active p38 MAP kinase inhibitor, for the potential treatment of inflammation, rheumatoid arthritis and cardiovascular diseases [7].
- SB590885 (associated with *BRAF* mutation): SB590885 is used for treating melanoma. SB590885 is a potent and selective ATP competitive inhibitor of B-Raf kinase [31].
- LFM-A13 (associated with *ANK2* amplification and *CELSR1* amplification and deletion): LFM- A13 is a potent and selective inhibitor of Bruton’s tyrosine kinase (BTK) used as an anti-breast cancer drug [2].
- GW-441756 (associated with *SYNE2* amplification): GW-441756 is a potent, selective inhibitor of the NGF receptor tyrosine kinase A (TrkA) [33].
- GNF-2 (associated with *PPFIA4* mutation): GNF-2 inhibits the cellular tyrosine phosphorylation of BCR-ABL in a dose-dependent manner induces a significant decrease in the levels of phospho-Stat5 in Ba/F3.p210 cells. [1].
- TW-37 (associated with *ITGB4* gene copy number): TW-37 is used for pancreatic cancer treatment. It is a small-molecule inhibitor of Bcl-2, inhibits cell growth and induces apoptosis in pancreatic cancer [28].
- WH.4.023 (associated with *PPFIA4* mutation): We do not have enough online data for this drug.
- Motesanib (associated with *PDGFRA* copy number): Motesanib is used for treatment of non-small cell lung cancer. an inhibitor of VEGR, cKit and PDGF receptor tyrosine kinases [24].

Detailed results for all the annotated drugs are available in additional file S2. To get deep into our analysis, we looked closely at the interaction between different variants, genomic expression, copy numbers, amplifications, and deletions with drug response. Figure 2 and 3 summarize the significant association and interaction between different genomic features and a particular drug.

**Figure 2:**
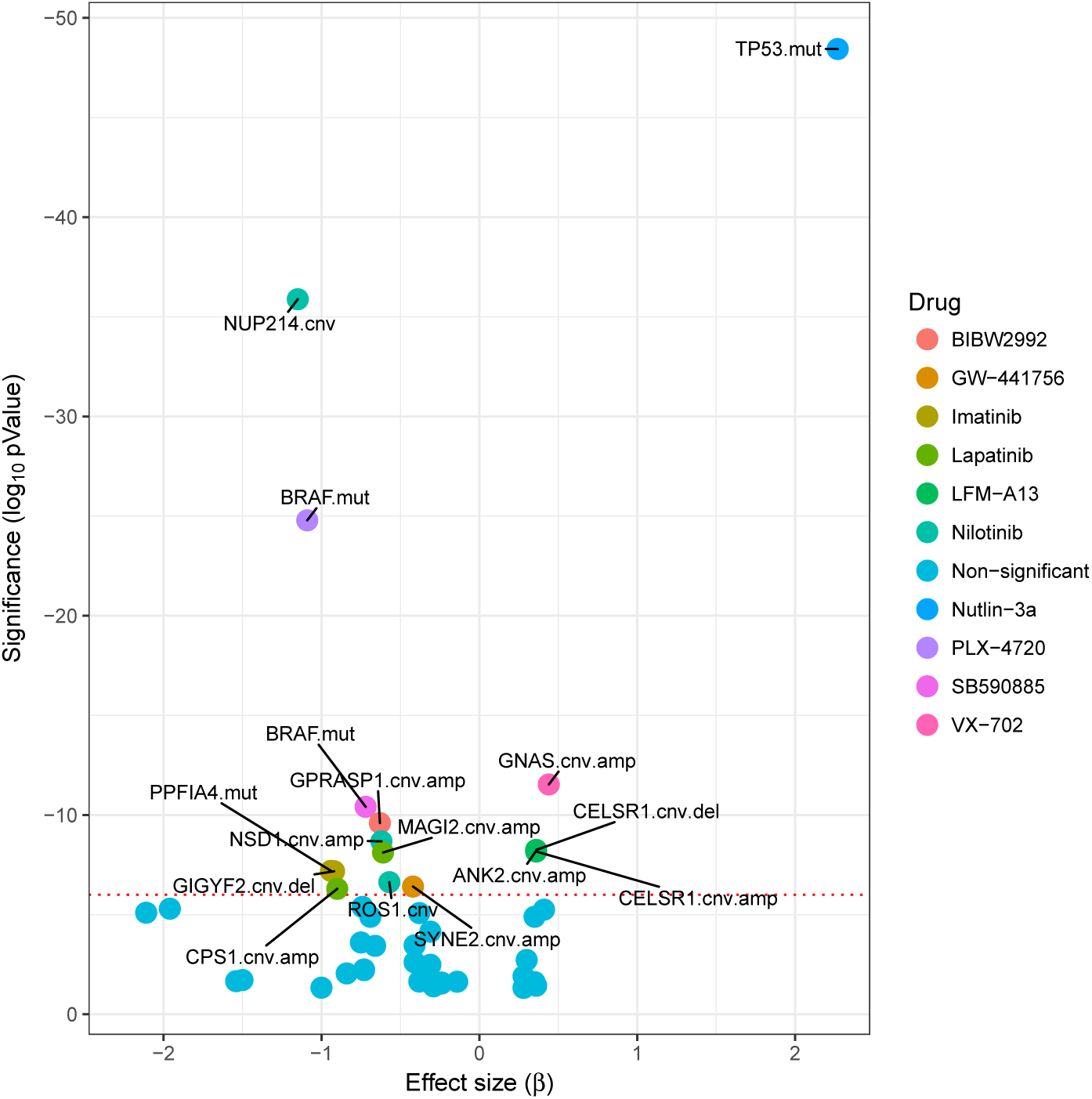
Significant association between different genomic features and a particular drug.

**Figure 3:**
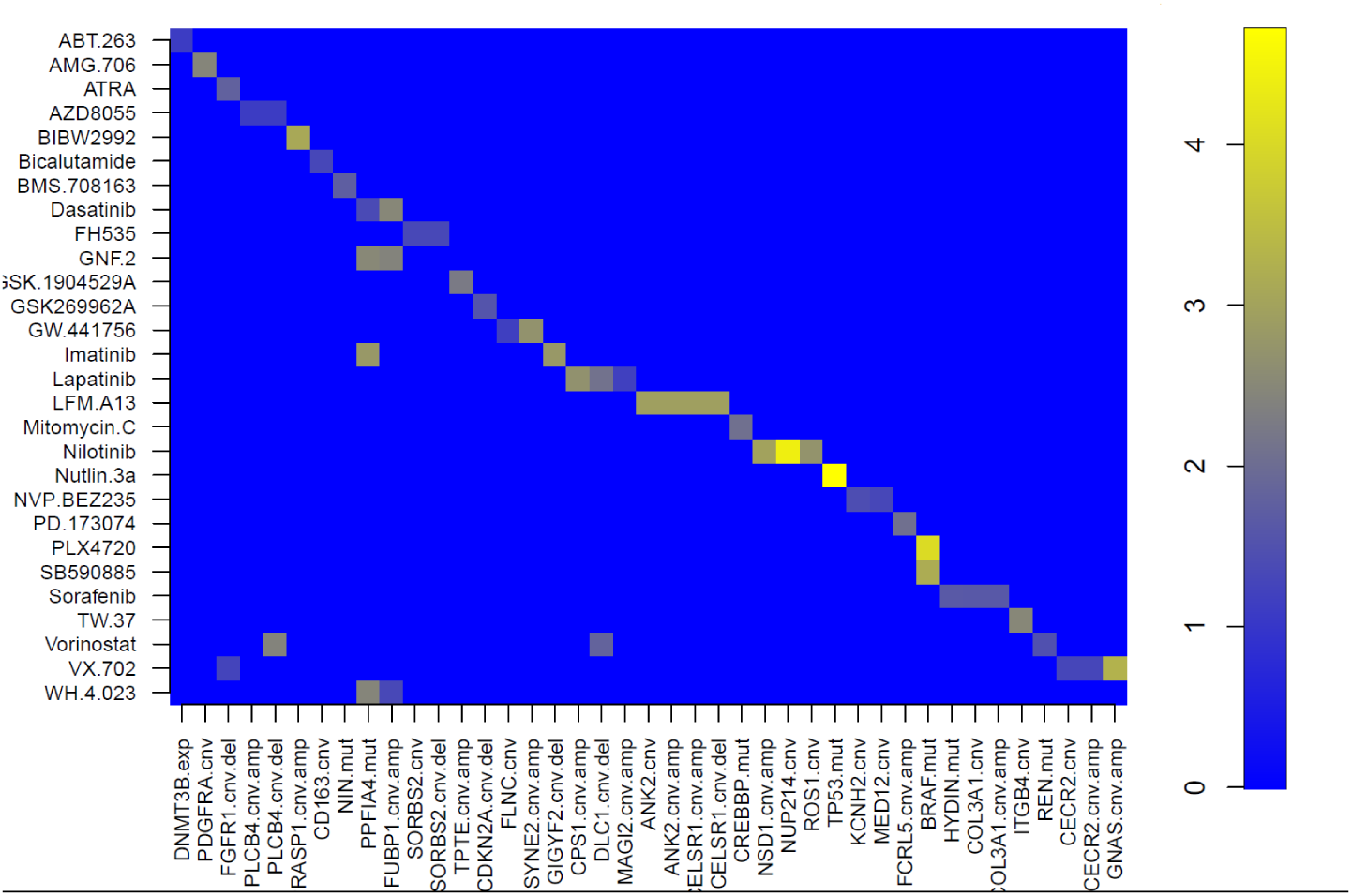
Summarizes the significant interaction between different genomic feature and a particular drug. For example one can see that genes *ANK2* copy number variation, and *CELSER1* copy number variation and *CELSR1* amplification are interacting with each other and are highly associated as a group with LFM-A13.

From figure 3 we can see that gene *ANK2* copy number variation, gene *CELSER1* copy number variation and amplification are highly associated with LFM-A13. We also find that gene *NSD1* amplification, gene *NUP214* copy number and gene *ROS1* copy number are as a group highly associated with Nilotinib. Finally, we find that gene *BRAF* mutation is interacting with the BRAF-selective inhibitors PLX4720 and SB590885.

## 3 Materials and Methods

We have screened and integrated a panel of cancer cell lines including mutation, DNA copy number, and gene expression data along with their responses to targeted and cytotoxic drug therapies from different databases. The dataset contained a broad range of drugs (5 categories) which are: targeted agents, cytotoxic chemotherapeutics, approved drugs used in clinical practice, drugs in development undergoing studies in clinical trials and experimental compounds.

### 3.1 Multi-dimensional genomic data

The data was collated together primarily from the publicly available GDSC data (version 5) [34], and IC50 measurements for a large number of drugs, against a wide range of cell lines [9].

### 3.2 Construction of genetic alteration profiles of candidate genes

Complete gene expression, mutation and copy number variation (cnv) data was downloaded from the publicly available Catalogue of Somatic Mutations in Cancer (COSMIC) dataset [8]. Mutation data was represented as a binary variable for each gene as mutated/non-mutated. Binary amplification and deletion variables were derived from copy number variation as cnv ≥ 5, and cnv *<* 2 respectively. A continuous value column, Δcnv, was also included representing amplifications as positive number and deletions as negative numbers to account for the magnitude of copy number variation.

The Broad Institute’s MutSig program [17] was used to identify commonly mutated genes in the publicly available data downloaded from The Cancer Genome Atlas (TCGA). The most commonly mutated 100 genes in the TCGA list were collected. To this list were added the 1000 most commonly mutated genes in the COSMIC dataset, giving a list of 1092 unique genes for analysis. The final data for the analysis contained the mutation status, copy number variation and expression data for these genes, giving a final dataset of 689 cell lines for which we have IC50 values for 141 drugs, mutation statuses for 1091 genes, copy number variation data for 476 genes, and expression data for 1025 genes. The final dataset, containing mutation, cnv and expression data, therefore has 3544 features. All data was normalized as explained next.

### 3.3 Processing each data set

Data set was first curated to include the samples of interest and all the information for each sample that the algorithm will use, such as drug subtype and cell lines types. Expression values in each data set are normalized and scaled by subtracting the mean and dividing by two standard deviations as in [10]. Categorical variables are binarized. Mutation and copy number variables have been centered by subtracting their mean. Missing expression values (for genes whose expression is present for some samples and not others in the data set) are imputed using nearest neighbor imputation (“knnImputation” in R package) [26].

### 3.4 Statistical analysis using bastah

An essential component of gathering all the information from different databases as explained is the systematic integration of large-scale genomic and drug sensitivity datasets. To identify genomic markers of drug response, we currently use a comprehensive analytical approach bastah whose goal is to correlate drug sensitivity (IC50 values and slope of the dose/response curve) with genomic alterations in cancer including point mutations, amplifications and deletions of common cancer genes for each drug. The approaches identify individual genomic features associated with drug sensitivity, and for each drug-gene association reports an unbiased size effect and statistical significance of the association with an adjusted p-value as will be briefly explained in this section.

In a linear regression model, we are given *n* pairs (*y*_1_; **x**_1_), (*y*_2_; **x**_2_)*,….,* (*y*_*n*_; **x**_*n*_), with vectors **x**_*i*_ *∈ ℝ*^*p*^ (*p*=3544) and response variables *y*_*i*_ given by

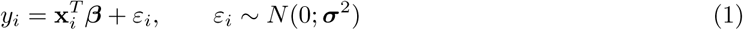

where ***ε*** is the error (noise) term with independent and identically distributed (i.i.d.) components having *E*[***ε***] = 0 and *V ar*(***ε***) = ***σ***^2^. An intercept term may be implicitly present. In matrix form, letting **y** = (*y*_1_, ….*,y*_*n*_)^*T*^ and denoting by **X** the design matrix with rows 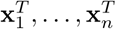,we have

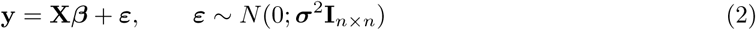

The goal is to estimate the unknown vector of parameters ***β ∈ ℝ****p*.

In the high-dimensional setting where *p>> n*, the matrix (**X**^*T*^ **X**) is rank deficient and one has to resort to biased estimators. A particularly successful approach is the lasso [25] which promotes sparse reconstructions through an ℓ_1_ penalization:

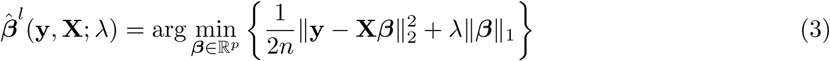

where *λ* is the tuning parameter that determines the amount of penalization. A larger value of *λ* tends to encourage a greater number of the *β*_*j*_ to be set exactly to zero. The optimal value of *λ* can be determined by cross validation on regression error, or via an information-theoretic test based on Bayesian Information Criteria.

Using [29] and [14], we can furthermore establish a distribution of 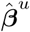 (an unbiased *β* estimator) that is

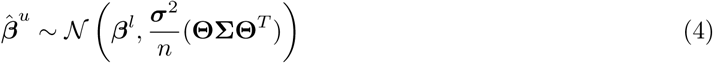

From which we derive the hypothesis test about ***β*** to test the null hypothesis *H*_0_ : *β*_*j*_ = 0 versus the alternative *H*_1_ : *β*_*j*_*≠*0 (two sided test statistics)

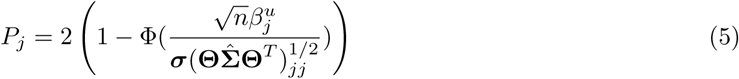

### 3.5 Model fitting procedures

The genomic features matrix **X** contains information about gene mutations, gene copy number variation, and gene expression characteristics for each cell line. The rows of the matrix correspond to the cell lines, there are 689 cell lines, and each column represent a specific genomic feature. There are 3544 genomic features columns in total. These columns are divided into 1091 gene mutation, 1428 gene copy number and 1025 gene expression.

The gene mutations columns in the genomic features matrix are binary, representing two categories (0 →no mutation in gene, 1→gene contains mutation). The gene copy number columns are divided into three types for each gene: gene.cnv, gene amplification (gene.cnv.amp), and gene deletions (gene.cnv.del). The gene.cnv columns are continuous (values *<* 0 are deletions, values≥3 are amplifications), while the gene amplification and deletion columns are binary (0→no amplification/deletion in gene, 1→gene is amplified/deleted). The last group of columns of the genomic features matrix is the gene expression columns. The expression values are continuous. Each column contains the expression values of a specific gene across all cancer cell lines. The last group of columns of the genomic features matrix is the gene expression columns. The expression values are continuous. Each column contains the expression values of a specific gene across all cancer cell lines

This TCGA and COSMIC input dataset was divided into five non-overlapping sample groups, used as cross-validation folds for training and testing data. For each cross-validation fold, each model was trained on 4/5ths of the samples, and used to make predictions of sensitivity for the held out 1/5th of samples. Within each training step, a separate 5-fold cross-validation procedure was employed for parameter tuning of each model.

### 3.6 Feature selection

We use our approach of sparsity inducing regularizers embedded within the estimation of linear regression,bastah described before to achieve feature selection. In implementing l_1_-regularized regression, the choice of regularization term is crucial. This procedure was repeated 100 times for each drug in order to assess the stability of the features (as done by [9]) when applying the 10-fold cross validation procedure. For each of the 100 runs, a feature list was built for the drug comprised of different genomic features with weights assigned to each. The final signature of markers for a drug consisted of all features that appear in any of the 100 runs along with significant p-value. The weights (w) and their respective p-values were used to assess effect sizes of features in a drug as a marker signature. The effect size of a feature was calculated by multiplying the feature’s weight by the p-values across the cell line panel. The effect size is therefore a normalization of the feature’s weight to account for the different scales used to measure the different genomic features.

Features with higher stability of correlation in cross-validation were considered of the highest confidence of truly being associated with drug response. The most significant features associated with drug response are those with large frequency and small p-value. In total, 40 features were highly reproduced and significantly associated with IC50 for 139 drugs. It is noticeable that copy number variation features have the wider ranges of average coefficients compared to gene mutation and expression features (Figure 4).

**Figure 4:**
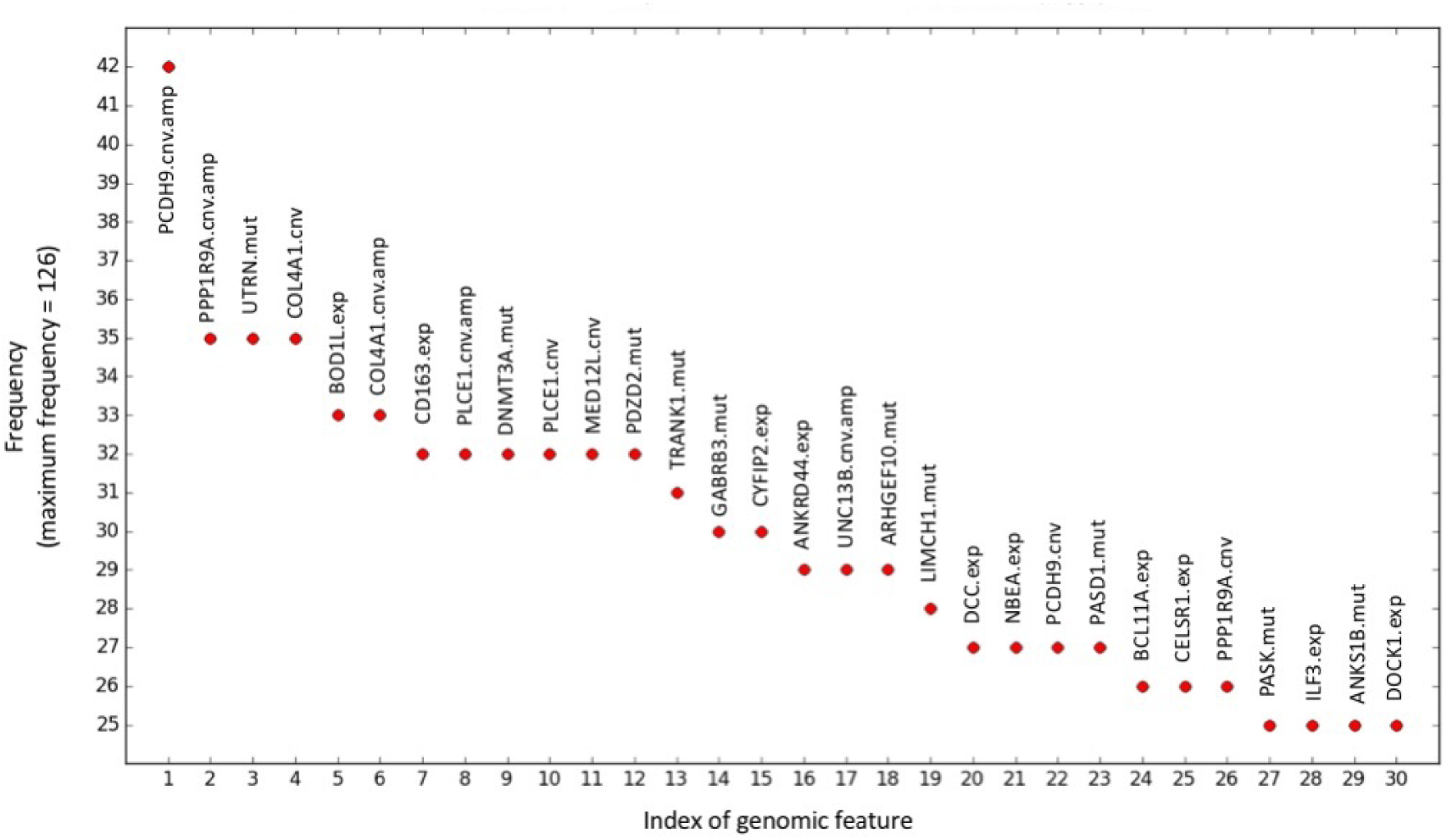
Top 30 most common significant genomic features among the panel of drugs.

## 4 Discussion

We have shown that models constructed using variants, genomic expression, copy numbers, amplifications, deletions and drug IC50 values, from a large panel of cell lines, can predict the chemotherapeuticresponse in patients. While acquired resistance to chemotherapeutics or targeted agents is well recognized and is the subject of recently intensive studies [19, 15], the variation in sensitivities to these drugs displayed by the cell lines in culture is not likely to be due to resistance acquired from previous exposure to these drugs; the patients from whose tumors the cell lines were derived would not have been treated with many of these drugs.

Variation of gene expression levels in cancer is known to be brought about by copy number variation, deletion and amplification and in this way the three variables ought to be considered simultaneously when scanning for driver genes. We likewise considered cancer driver gene selection by means of investigation of copy number, mutation, amplification, deletion and expression levels [23] via newly developed generalized lasso techniques. When using gene network analysis on gene expressions only (results we did not report here), no structure was found associating drug sensitivity to gene expression. When variation in gene expression levels was used with penalized lasso, we obtained more information.

It might appear to be astounding that quality and variation in gene expression alone has such exceptional influence in enhancing drug responses, as the approach disregards what are thought to be important factors in drug sensitivity, for example, malignancy type or other specific markers. We demonstrated, through this work, that the good performance may (partially) come from not only the way the unbiased test statistics were used for genomics features prediction but also that the variability in gene expression as a surrogate for unmeasured phenotypes are straightforwardly important to chemotherapeutic sensitivity, therefore, can catch parts of both germline and tumor-specific genome variation.

In summary, we have proposed a novel integrative computational strategy based on a multidimensional multitask penalized regression for identifying driver genes for cancer. To effectively perform high dimensional genomic data analysis, we used recursive strategies in line with the penalized lasso method. Furthermore, we have proposed a parametric multidimensional statistical test for gene selection based on robust regression modeling. These findings have profound implications for personalized medicine and drug development. Future work will focus on improving predictions using more rigorous transcriptome quantification.

